# Chromosome-scale genome assembly of *Fusarium oxysporum* strain Fo47, a fungal endophyte and biocontrol agent

**DOI:** 10.1101/2020.05.22.110718

**Authors:** Bo Wang, Houlin Yu, Yanyan Jia, Quanbin Dong, Christian Steinberg, Claude Alabouvette, Veronique Edel-Hermann, H. Corby Kistler, Kai Ye, Li-Jun Ma, Li Guo

## Abstract

Here, we report a chromosome-level genome assembly of *Fusarium oxysporum* strain Fo47 (12 pseudomolecules; contig N50: 4.52Mb), generated using a combination of PacBio long-read, Illumina pair-ended and Hi-C sequencing data. Although *F. oxysporum* causes vascular wilt to over 100 plant species, the strain Fo47 is classified as an endophyte and widely used as a biocontrol agent for plant disease control. The Fo47 genome carries a single accessory chromosome of 4.23 Mb, compared to the reference genome of *F. oxysporum* f.sp. *lycopersici* strain Fol4287. The high-quality assembly and annotation of the Fo47 genome will be a valuable resource for studying the mechanisms underlying the endophytic interactions between *F. oxysporum* and plants, as well as deciphering the genome evolution of the *F. oxysporum* species complex.

## Genome announcement

A filamentous ascomycete fungus, *Fusarium oxysporum* represents a species complex of diverse members that cause soil-borne vascular wilt diseases to over 100 plant species in a host-specific fashion (Kistler et al. 1997; Ma et al. 2013; Edel-Hermann and Lecomte 2019). Strains pathogenic to specific host species are classified as *formae speciales* (f. sp.), and, together with related non-pathogenic strains, they are collectively known as the *F. oxysporum* species complex (FOSC) (Kistler et al. 1997; Michielse and Rep 2009). Although notorious as plant pathogens, commonly *F. oxysporum* strains are plant endophytes often beneficial to host plants (Alabouvette et al. 2009). Fo47, a strain isolated from plant disease suppressive soils (Alabouvette 1986), grows endophytically and is widely used as biological control agent for preventing vascular wilt of tomato caused by *F. oxysporum* f.sp. *lycopersici* (Fol) (Alabouvette et al. 2009) and other plant diseases as well (He et al. 2002; Paparu et al. 2007; Veloso and Díaz 2012). In general, beneficial microbes contribute to the fitness of host plants potentially through diverse mechanisms including induced systemic resistance, promoting nutrient assimilation and conferring antagonisms against microbial pathogens (Alabouvette et al. 2009; Van der Ent et al. 2009; Aimé et al. 2013; Pieterse et al. 2014). The biocontrol effect of Fo47 is widely exploited in agriculture but the molecular mechanisms behind it remain poorly understood. Decoding the genome of Fo47 will provide a valuable resource for not only facilitating dissection of such mechanisms, but also contributing to comparative genomic studies for understanding the genome evolution of FOSC in terms of endophytism, pathogenesis and host specificity. The genome assembly of the same strain was previously deposited at NCBI under the accession number GCA_000271705.2.

High molecular weight genomic DNA was extracted from four days old Fo47 mycelia using CATB (Cetyl Trimethyl Ammonium Bromide) method. A total of 6.85 Gb paired-end short sequence reads (2×150bp) and 18.09 Gb single-molecule long-read sequence reads were generated using Illumina NovaSeq 6000 platform and PacBio SMRT RS II platform, respectively. K-mer (17mer) frequency analysis of 6.85 Gb Illumina reads (138X coverage) using Jellyfish-GenomeScope pipeline (Marcais and Kingsford 2011; Vurture et al. 2017) revealed an estimated genome size of 49.54 Mb for Fo47. The 18.09 Gb (341X coverage) PacBio SMRT data (1.24 million reads) has a read length N50 of 18,156 bp after filtering. We then assembled Fo47 genome from the PacBio reads using the assembler *NextDenovo* v2.0 (https://github.com/Nextomics/NextDenovo). The PacBio assembly was further polished by Illumina PE reads using *Pilon* v. 1.22 (Walker et al. 2014) to correct 2,040 (0.004%) erroneous positions. To make the chromosome-scale assembly, the contigs were scaffolded using chromatin contact maps produced by Hi-C (high throughput chromosome conformation capture) reads. Basically, a single Hi-C library prepared from cross-linked chromatins of fungal cells using a standard Hi-C protocol (Belton and Dekker 2015) were sequenced by NovaSeq 6000 to yield 5.76Gb (115x coverage) Illumina PE reads. The Hi-C data was used to anchor all contigs on 12 pseudomolecules (chromosomes) using Juicer v. 1.5 (Durand et al. 2016a) and Juicebox v. 1.11.08 (Durand et al. 2016b), followed by manual checks. The number of chromosomes was determined by centromeric interaction regions detected in Hi-C contacted map figures (Marbouty et al. 2014).

The final assembly of the strain Fo47 genome is 50.36 Mb placed in 12 chromosomes, proximate to its estimated genome size. Each chromosome is represented in a single contig, with a contig N50 of 4.52Mb, demonstrating the high continuity of this assembly (Table 1). Out of 12 chromosomes, nine had more than three copies of the telomeric repeat unit TTAGGG on both ends, whereas the other three contained those copies on only one end, indicating a high-quality Fo47 genome assembly of chromosome level. Compared to a previous Fo47 genome assembly (GenBank accession: GCA_000271705.2) at NCBI (Table 1), the major improvement happens at chromosome Chr7, as 73 contigs (3.96Mb) of the initial assembly can be aligned to the single 4.23Mb Chr7 of the new assembly using *Mummer* v. 3.1 (Kurtz et al. 2004).

**Table 1.** Summary of the genome assembly and annotation statistics of *Fusarium oxysporum* Fo47 compared to a previous assembly of Fo47 (GenBank accession: GCA_000271705.2) and *F. oxysporum* f. sp. *lycopersici* Fol4287 (GenBank accession: GCA_000149955.2) (Ma *et al*. 2010).

| Strains | Fo47 (this study) | Fo47 (GCA_000271705.2) | Fol4287<sup>b</sup> |
| --- | --- | --- | --- |
| Number of contigs | 12 | 124 | 88 |
| Number of chromosomes | 12<sup>a</sup> | NA<sup>c</sup> | 15 |
| Number of lineage-specific chromosomes | 1 | NA<sup>c</sup> | 4 |
| Genome size (Mb) | 50.36 | 49.67 | 61.47 |
| GC content (%) | 47.71 | 47.68 | 48.48 |
| Contig N50 (bp) | 4,523,040 | 3,844,136 | 4,589,937 |
| Number of protein coding genes<sup>d</sup> | 17,234 | 17,055 | 17,637 |
| Repeat content (%) | 3.42 | 2.77 | 5.43 |
| Genome assembly completeness (%) <sup>e</sup> | 98 | 96 | 99 |
| Predicted proteome completeness (%) <sup>e</sup> | 99 | 99 | 98 |

a. Number of putative centromeres was determined using Hi-C contact map.
b. Reported by Ma *et al.* (2010).
c. NA: unavailable
d. Protein-coding genes were identified by the same pipeline (this study) to ensure a fair comparison.
e. Completeness = Total BUSCOs - Missing BUSCOs / Total BUSCOs \* 100

For genome annotation, *GeneMark-ES* (Borodovsky and Lomsadze 2011) was first used for *de novo* gene structure prediction. A *Fusarium*-specific gene model was then used to train *Augustus* v. 3.1 (Mario and Burkhard 2005). *MAKER2* pipeline (Cantarel et al. 2008) was used to find protein-coding genes integrating gene models predicted from *GeneMark-E*S and *Augustus* and protein sequences of the *F. oxysporum* reference genome of a tomato pathogenic isolate Fol4287 (GenBank assembly accession: GCA_000149955.2) (Ma et al. 2010), with RepeatMasker v. 4.0.7 (Tarailo-Graovac and Chen 2009) option on to find and mask repetitive elements. In total, the Fo47 genome has 17,234 protein coding genes and 3.42% repeat content (Table 1). The genome assembly completeness was assessed by BUSCO (Benchmarking Universal Single Copy Ortholog) (Simão *et al*. 2015) using both genome and proteome sequences, showing BUSCO score of 98% (genome) and 99% (proteome) based on the ‘fungi_odb9’ library (Table 1), suggesting a high degree of completeness for this Fo47 genome assembly.

Compared to the reference genome Fol4287 (Ma et al. 2010), Fo47 has a smaller genome size, fewer genes and repeat elements (Table 1). A whole genome sequence alignment between Fo47 and Fol4287 using *Mummer* v. 3.1 (Kurtz et al. 2004) reveals that the size of sequence mapped to core genome of Fol4287 (11 chromosomes) (Ma et al. 2010) was 46.10Mb, suggesting a set of 11 core chromosomes in Fo47. A single Fo47 chromosome Chr7 (4.23Mb) did not align to any Fol4287 chromosome, indicative of a lineage-specific chromosome in Fo47. Chr7 is highly enriched for transposable elements (TE), with 51% of total TE sequences located on Chr7 and 49% located on the rest of genome. Over representation of repetitive sequences in this chromosome explains the fragmentation in the initial assembly. Chr7 has a smaller average gene density (290 per Mb) compared to core chromosomes (347 per Mb). The 1,247 genes on Chr7 are significantly enriched (P-value: 1.5e-4) for regulators of G-protein signaling functions suggesting their likely role in external sensing and organism response to external nutrient sources and other environmental signals (Bayram and Braus 2012; Kim et al. 2017). This genome assembly of Fo47 will be a useful resource for comparative genomics of *F. oxysporum*, and future functional genomic studies of Fo47 and host endophytic interactions. The assembled genome and gene annotations have been deposited at GenBank under the accession numbers CP052038-CP052049. PacBio reads, Illumina reads and Hi-C Illumina data are available in the NCBI Sequenced Read Archive under the accession numbers SRR11526760, SRR11523115, and SRR11523120, respectively.

## Acknowledgment

We would like to thank the anonymous reviewers for their kind and helpful comments on the original manuscript.

## Funding

This project was supported by the National Natural Science Foundation of China (31701739, 31970317), Fundamental Research Fund of Xi’an Jiaotong University, and National Key R&D Program of China (2018YFC0910400).

## Notes

### Competing Interest Statement

The authors have declared no competing interest.

